# HESTA: a curated and reusable database for the human early organogenesis spatiotemporal transcriptome atlas

**DOI:** 10.64898/2026.05.28.728391

**Authors:** Zhicheng Xu, Yuejiao Li, Weiwen Wang, Ying Zhang, Lidan Fan, Jing Chen, Wensi Du, Tao Yang, Ya Gao, Kailong Ma

**Author notes:** These authors contributed equally (Zhicheng Xu, Yuejiao Li, Weiwen Wang). To whom correspondence should be addressed:Kailong Ma[ ].

## Abstract

**Background:** Human organogenesis is orchestrated by precise spatiotemporal gene expression. Mapping these dynamic processes requires transcriptomic data that preserve native anatomical context across continuous developmental stages.

**Findings:** We present a spatiotemporal transcriptome database of human embryogenesis, profiling 77 sagittal sections from 13 euploid embryos (CS12–CS23) using Stereo-seq, yielding 14,708,858 bin50 spots. The atlas annotates 50 organs and maps 198 molecularly distinct substructures, complemented by 607,093 snRNA-seq cells. The database features a Spatial Exploration module for locating sections and visualizing spatial distributions of organs and substructures, and an Organ Atlas module for visualizing gene expression, regulon activities, and pathway enrichment at the single-organ level across stages.

**Conclusions:** This database provides an interactive resource to access spatial gene expression, substructures, and regulatory networks across 50 developing human organs, supporting further research into the mechanisms of human organogenesis.

## Data Description

### Context

Human embryonic development is coordinated by precisely regulated spatiotemporal gene expression programs. Following gastrulation, the embryo enters a critical period of early organogenesis (1-4), during which major organ primordia are established, cellular diversity expands rapidly, and tissue architecture becomes increasingly complex. Carnegie stages (5) 12–23, corresponding approximately to post-conception weeks 4–8, represent a particularly important developmental window characterized by extensive diversification of cell types and dynamic organ morphogenesis. During this period, the embryo is especially vulnerable to teratogenic perturbations. Developmental disturbances during embryogenesis have been associated with birth defects (6) and neurodevelopmental disorders(7).

Single-cell and single-nucleus transcriptomic studies have provided important insights into cellular heterogeneity during human development. However, organogenesis is not solely determined by cell identity, it also depends on the spatial organization of cells, anatomical context, and interactions with neighbouring tissues. Spatial transcriptomic technologies therefore provide an essential layer of information by preserving the physical location of gene expression signals within developing tissues. For human early organogenesis, such spatially resolved data are especially valuable, as many organs are small, rapidly changing, and tightly folded, making it difficult to interpret gene expression without anatomical context.

Recently, a whole-embryo spatiotemporal transcriptomic atlas of post-gastrulation human embryos was published, covering 13 euploid embryos and 77 sagittal Stereo-seq sections from CS12 to CS23. This atlas annotated 50 organs or anatomical regions and 198 molecularly defined spatial substructures, and integrated single-nucleus RNA sequencing data to provide cell-type-level references for spatial interpretation(8). Together, these data provide a comprehensive framework for studying organ-specific differentiation, spatial regulatory programs, developmental trajectories, and allelic expression patterns during human early organogenesis.

Although the original atlas study establishes the biological value of the atlas, the resulting datasets are large, multidimensional, and difficult to reuse directly. Users seeking to explore gene expression across developmental stages, compare spatial patterns among sections, inspect organ or substructure annotations, evaluate pathway or regulon activity, or download processed matrices for downstream analysis would otherwise need to navigate multiple files and reconstruct complex relationships among embryos, stages, sections, organs, substructures, cell types, and derived data products.

To address this need, we developed HESTA, the Human Embryo Spatiotemporal Transcriptomic Atlas, as a curated and reusable data resource for the spatiotemporal transcriptomic atlas of human early organogenesis (https://db.cngb.org/hesta/; Figure 1). HESTA organizes the companion atlas into standardized metadata, section-level spatial matrices, organ and substructure annotations, single-cell reference data, selected organ-level gene, pathway and regulon matrices, allele-specific expression maps, three-dimensional reconstruction data, and interactive web interfaces for browsing, querying, downloading and reuse. Rather than serving only as a visualization website, HESTA provides a data access layer that connects atlas-level annotations with processed and derived data products. A summary of the data scale across different developmental stages is provided in Table 1.

**Table 1.**
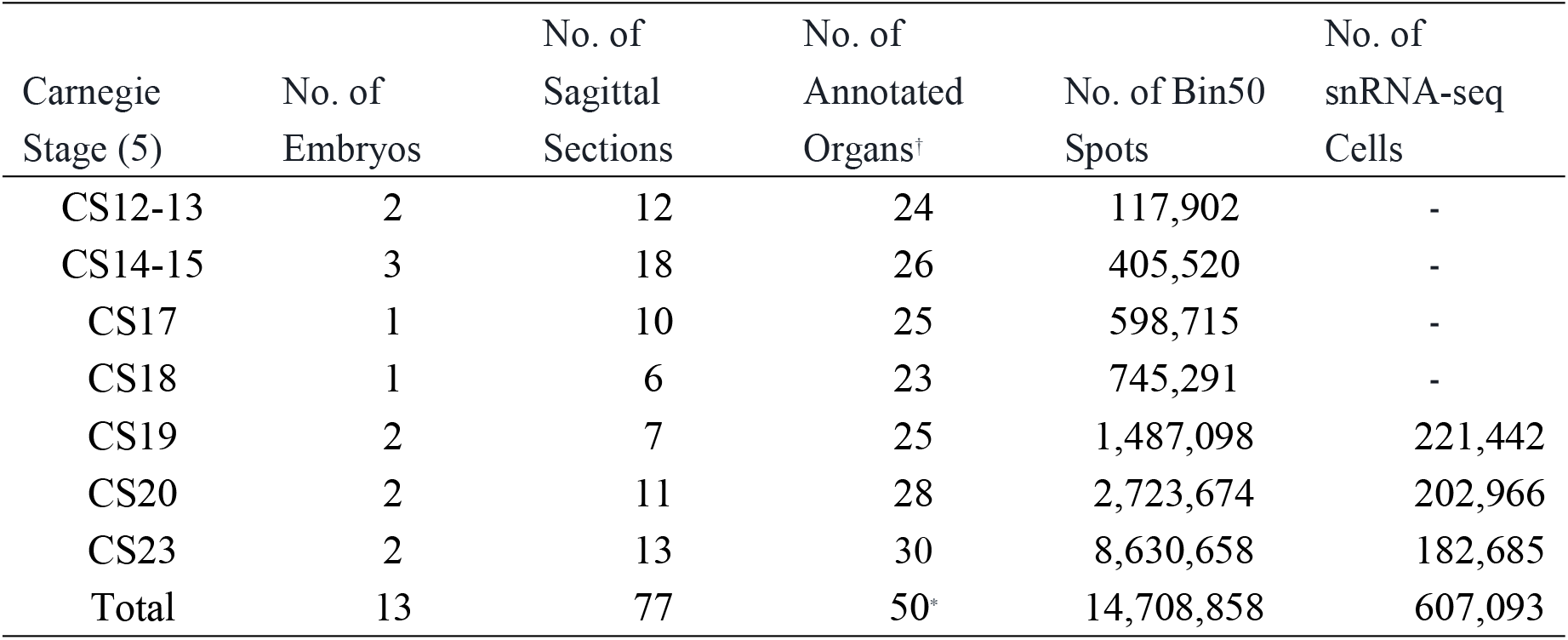
Summary of HESTA data resources across Carnegie stages (CS12–CS23). For each developmental stage, the table reports the number of profiled euploid embryos, sagittal Stereo-seq sections, anatomically annotated organs or regions, bin50 spatial spots, and single-nucleus RNA sequencing (snRNA-seq) cells. snRNA-seq data are available only for selected stages (CS19, CS20, and CS23) as generated in the original atlas study. The “Total” row summarizes the cumulative data scale across all stages. †The count of embryonic organs excludes “Extraembryonic Tissue” and “Cavity”, which are retained in the h5ad matrices for spatial completeness but are not anatomically defined organs. *The total count of 50 represents unique organs across all stages rather than a simple sum of per-stage counts.

| Carnegie Stage (5) | No. of Embryos | No. of Sagittal Sections | No. of Annotated Organs<sup>†</sup> | No. of Bin50 Spots | No. of snRNA-seq Cells |
| --- | --- | --- | --- | --- | --- |
| CS12-13 | 2 | 12 | 24 | 117,902 | - |
| CS14-15 | 3 | 18 | 26 | 405,520 | - |
| CS17 | 1 | 10 | 25 | 598,715 | - |
| CS18 | 1 | 6 | 23 | 745,291 | - |
| CS19 | 2 | 7 | 25 | 1,487,098 | 221,442 |
| CS20 | 2 | 11 | 28 | 2,723,674 | 202,966 |
| CS23 | 2 | 13 | 30 | 8,630,658 | 182,685 |
| Total | 13 | 77 | 50* | 14,708,858 | 607,093 |

**Figure 1.**
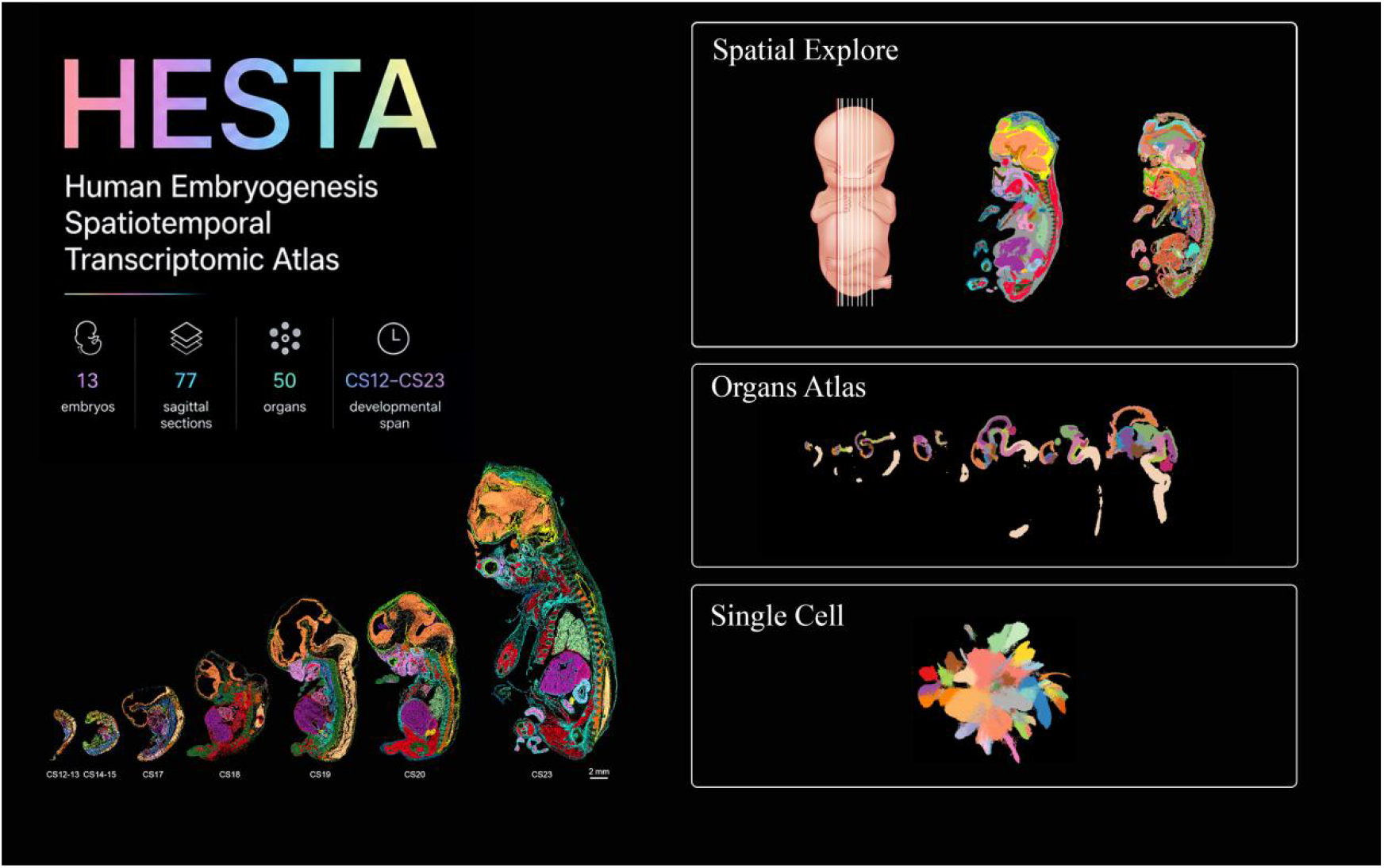
Overview of HESTA database. The database consists of three visualization modules: the Spatial Explorer for section-level organ and substructure annotations, the Organ Atlas for organ-based gene expression, pathway and regulatory networks, and the Single-cell module for reference mapping.

## Methods

### HESTA data catalogue and release overview

The HESTA data resource was organized around multiple biological and computational layers from 13 euploid human embryos across CS12–CS23. Section-level spatial matrices serve as the foundational data layer for whole-section visualization and spatial gene expression exploration. Substructure-level matrices were generated for representative sections with high-resolution substructure annotations. Single-nucleus RNA-seq data were incorporated as a reference layer for cell-type interpretation. Organ-level derived matrices were generated for selected organs and anatomical regions, including gene expression, pathway activity and regulon activity matrices.

Additional derived resources include allele-specific expression maps, three-dimensional reconstruction data and a Carnegie stage to post-conceptional week mapping table. These data products represent different biological and computational levels and therefore do not have uniform coverage across organs, sections or substructures.

### Processing and curation of section-level spatial matrices and organ annotations

Section-level spatial matrices were generated from Stereo-seq bin50 count matrices. For each section, gene expression profiles and spatial coordinates were used to support spatially constrained clustering. Expression-based neighbourhood graphs and spatial neighbourhood graphs were constructed using Scanpy(v1.8.2) (9) and Squidpy (v1.2.0) (10), respectively, and then integrated for Leiden clustering. The resulting clusters were annotated at the organ or anatomical-region level based on highly variable genes, marker gene expression and anatomical context.

The processed section-level matrices were stored as h5ad files, and the organ annotations were recorded in the .obs attribute of AnnData, with the key of ‘celltype’. Each file preserves the bin50-level expression matrix, spatial coordinates, section metadata and organ-level annotation labels. These files provide the foundational data layer for HESTA and support both interactive visualization and downstream reuse. In the web interface, section-level matrices are connected to Spatial Explorer, allowing users to inspect tissue-level spatial clustering, anatomical annotation maps and gene expression overlays.

### Processing and curation of substructure-level matrices

Substructure-level matrices were generated for 26 of the 77 representative sections. These sections, comprising two sections from each of the 13 embryos, were selected for high-resolution anatomical annotation, with cavitary organ regions excluded. To identify fine spatial domains, we applied two distinct strategies based on organ complexity:

For major organs such as the brain, spinal cord, and heart, secondary spatial clustering was performed directly at the individual section level using spatial expression matrices. For other organs, bin50 data from the 26 representative sections, along with additional sections capturing the broadest organ coverage, were aggregated across specimens to improve the detection of developmental substructures. For these pooled matrices, raw counts were integrated using scvi-tools (v1.2.2) (11) to generate batch-corrected latent representations.

In both strategies, the resulting representations were used for neighbourhood graph construction, UMAP embedding, and Louvain clustering with a resolution of 1.0. Substructure labels were assigned using established region-specific and cell-type-defining markers. The resulting substructure-level data products were stored as h5ad files, with organ and substructure labels saved in the .obs attribute under the keys ‘organ’ and ‘substructure’.

### Processing and curation of the single-cell reference dataset

The single-cell reference dataset was derived from the snRNA-seq data generated in the original atlas study. It provides cell-type-level expression profiles and annotations that support interpretation of spatial gene expression patterns. The original atlas study reported snRNA-seq data from CS19, CS20 and CS23 embryos, yielding 607,093 cells and resolving 68 cell types through reference-guided annotation.

The processed single-cell matrix was stored as an h5ad file, with cell-type annotations and sample identities recorded in the .obs attribute under the keys ‘celltype’ and ‘sample’, respectively. This file was integrated into HESTA as a reference layer for marker validation and cell-type interpretation. In the web interface, the single-cell module enables users to examine whether genes observed in spatial sections are enriched in specific cell types, thereby helping interpret mixed spatial bins and supporting organ or substructure annotation.

### Processing of organ-level gene, pathway and regulon matrices

Organ-level derived matrices were generated to support organ-centered exploration of developmental dynamics. For each selected organ or anatomical region, representative spatial sections from all available developmental stages were merged or organized into organ-specific data products. These organ-level files provide derived views of the atlas and complement the section-level spatial matrices and atlas-level organ annotations.

The organ gene matrices provide the core spatial expression layer. Structured as bin-by-gene matrices at bin50 resolution, they capture per-bin expression profiles for a given organ. These matrices support organ-specific gene expression queries across developmental stages and are integrated into the Organs module of HESTA.

To generate pathway activity matrices, gene expression matrices were converted into functional activity profiles. Gene Ontology Biological Process activities were quantified at the bin50 level using rank-based gene set enrichment analysis. In the current implementation, irGSEA (v3.2.5) (12) with the UCell algorithm was used to calculate per-bin50 enrichment scores against Gene Ontology Biological Process gene sets. The resulting m bin50 by n pathways matrices were converted into h5ad files for visualization and reuse.

To generate regulon activity matrices, regulatory networks were inferred from organ-specific raw count matrices using Stereopy (v1.5.1) (13) with the KNN method, AUCell enrichment scores were calculated to quantify regulon activity per bin50, generating a bin-by-regulon matrix.

These AUC_matrix were converted into h5ad files and integrated into the Organs module.

### Processing of allele-specific expression maps

To illustrate the spatial divergence of imbalanced allelic expression, allele-specific expression maps were prepared as precomputed derived visualization resources based on all sections from three embryos: CS14–15 embryo 1 (E1), CS14–15 E3, and CS20 E2. These high-confidence imbalanced genes were identified by integrating spatial transcriptomics with embryonic and maternal WGS, with long-read sequencing used to improve haplotype phasing accuracy.

Embryonic heterozygous variants (Genotype Quality (GQ) > 10, Read Depth (DP) > 4) were phased and aggregated to the gene level, and allelic reads were quantified using samtools. Confident imbalanced genes were defined as those containing more than three variants, or a single variant supported by more than 300 reads, and imprinting genes in the Geneimprint Database. For each section, genes passing the allele-specific expression filtering criteria were linked to spatial visualization maps. Each allele-specific expression map is linked to a section ID, gene symbol, statistical result, source data version, and visualization file. The web interface uses a section-based selection workflow: users first select a section, after which the corresponding list of significant genes is displayed for spatial visualization.

### Three-dimensional reconstruction

Three-dimensional reconstruction was generated using eight consecutive Stereo-seq sections from the CS12–13 E2 embryo. Section-level spatial data from each slice were converted into coordinate-indexed expression matrices and encapsulated as AnnData objects, with organ annotations recorded in the .obs attribute under the key ‘annotations’. Manual coordinate transformations were applied to correct orientation and mirroring differences among sections, followed by coordinate translation to remove absolute positional offsets.

After orientation harmonization, STitch3D (v1.0.3) (14) was used to perform three-dimensional alignment and spatial registration across consecutive sections. The reconstructed data object was integrated into HESTA for interactive visualization, allowing users to inspect spatial relationships among embryonic structures in a three-dimensional context.

### HESTA web resource and software implementation

The web frontend was implemented using HTML, CSS and JavaScript. The backend was developed using Django, with Nginx serving as the reverse proxy server. Interactive visualization of spatial and single-cell datasets was powered by Cirrocumulus(15). The interface is organized into seven sections: Home, Spatial Explorer, Organ Atlas, Single-cell, Analysis, Download and Help.

## Data Validation and Quality Control

The spatial and single-nucleus transcriptomic datasets described here were first reported in the companion Research Article (8). The technical validity of these data is supported by rigorous quality control, including the removal of low-quality sections/cells via established thresholds, high cross-embryo and cross-section reproducibility, and concordance between Stereo-seq expression patterns and RNA *in situ* hybridization results for eight organ-specific markers. Furthermore, the biological resolution of these datasets has been demonstrated through their application to mapping developmental trajectories across 50 organs and 198 substructures, as well as to elucidating the spatial dynamics of organ-specific regulons and lncRNA-associated ontologies (8).

### Usecase:spatiotemporal visualization of organ-specific marker genes

Based on these curated and quality-controlled datasets, we next used HESTA to illustrate how users can explore spatially resolved gene expression programs during human early organogenesis. As an example, representative marker genes were queried across CS12-CS23 through the spatial visualization module (Figure 2). The resulting plots showed clear stage-dependent and anatomically restricted expression patterns, providing an intuitive view of how organ identities and tissue-specific transcriptional programs progressively emerge during embryonic development. In the whole-embryo sections, *FENDRR, ACTG2, SERPINA1*, and *FABP1* showed noticeably different spatial distributions, which broadly corresponded to their known developmental or organ-related functions (Figure 2A). *FENDRR* was mainly detected in areas associated with mesodermal development, particularly in gastrointestinal tract and lung. *ACTG2* showed enrichment in regions corresponding to developing visceral smooth muscle. This pattern became more apparent at later stages, when smooth muscle-associated structures are expected to become more organized. Given that *ACTG2* encodes an enteric smooth muscle actin, its expression pattern fits well with the maturation of gastrointestinal and other visceral smooth muscle compartments(16). By comparison, *SERPINA1* and *FABP1* displayed stronger signals in endoderm-derived organs, especially in regions corresponding to the developing liver and digestive tract. The consistent expression of *SERPINA1* in liver is in line with its well-established association with hepatic differentiation and secretory function(17). *FABP1*, a marker related to lipid metabolism and commonly enriched in liver and intestinal tissues(18), also showed a spatial pattern consistent with the emergence of metabolically active digestive organs. These observations suggest that HESTA can recover not only broad germ-layer-associated transcriptional programs, but also more specific organ-level maturation signatures.

**Figure 2.**
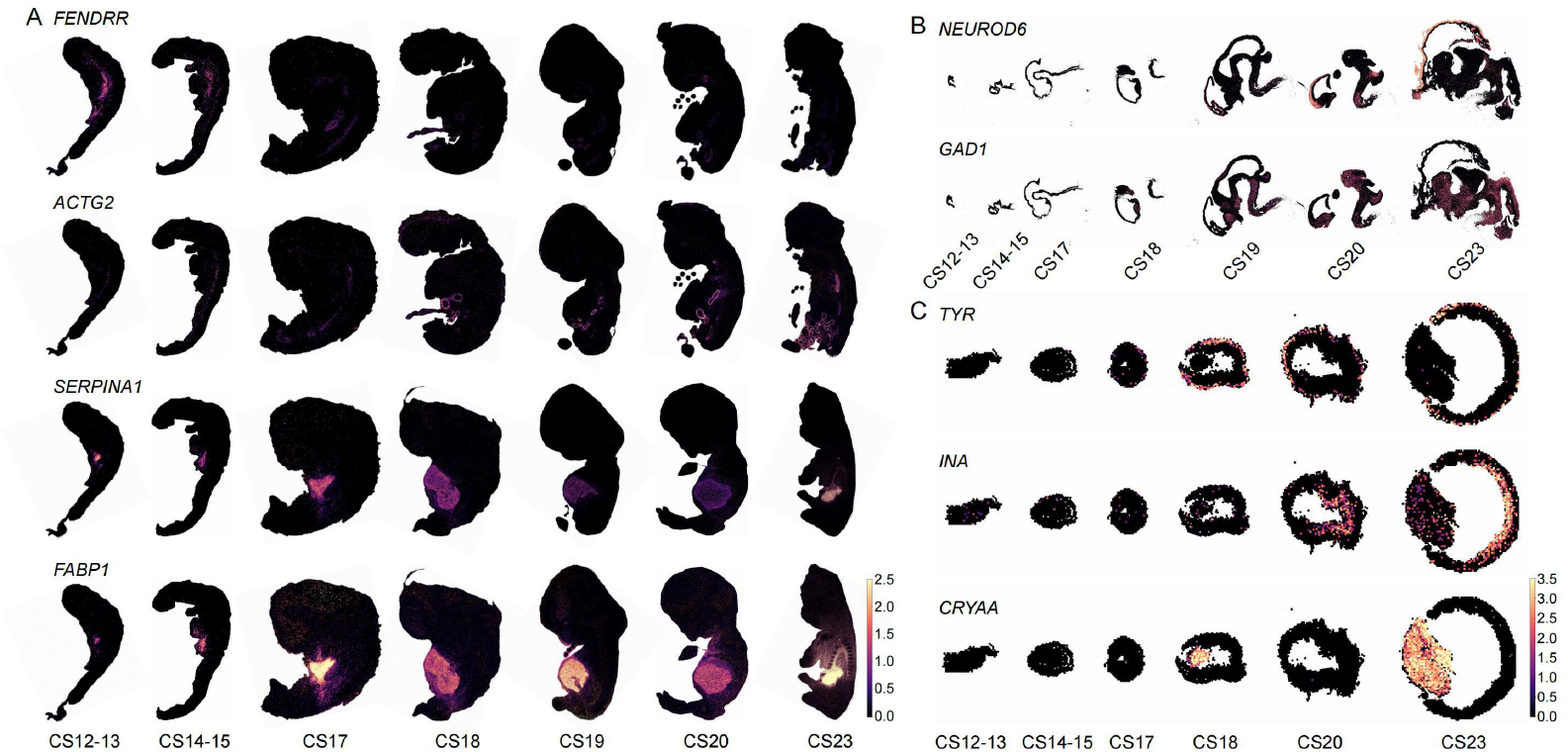
Spatiotemporal visualization of representative organ- and substructure-specific marker genes in HESTA. (A) Spatial expression of *FENDRR, ACTG2, SERPINA1*, and *FABP1* in whole-embryo sagittal sections across CS12-CS23. *FENDRR* marks mesoderm-associated regions, including the developing gastrointestinal tract and lung; *ACTG2* is enriched in visceral smooth muscle compartments; and *SERPINA1* and *FABP1* highlight endoderm-derived tissues, particularly the liver and digestive tract. (B) Substructure-level visualization of neural markers. *NEUROD6* is enriched in the pallidum, whereas *GAD1* is detected in the subpallium, midbrain, hypothalamus, and cerebellum, reflecting spatially distinct excitatory and inhibitory neuronal programs. (C) Spatial expression of eye-related markers. *TYR* is localized to melanocyte-associated ocular regions, *INA* is enriched in retinal ganglion cells, and *CRYAA* shows strong expression in the developing lens.

Beyond the organ-level analysis, users can also explore gene expression at the substructure level. *NEUROD6* and *GAD1* were mainly enriched in neural regions, with their expression becoming more spatially structured as development progressed (Figure 2B). *NEUROD6*, which is involved in the differentiation and maturation of excitatory neurons(19), showed particularly strong expression in the pallidum. *GAD1*, a key enzyme required for GABA synthesis(5), was detected in the subpallium, midbrain, hypothalamus, and cerebellum, marking regions where inhibitory neuronal circuits start to form. The eye-related markers showed particularly clear anatomical restriction, too (Figure 2C). *TYR* was detected in melanocytes, in line with its function as the rate-limiting enzyme for melanin biosynthesis in the retinal pigment epithelium. Because ocular melanin is important for retinal development, visual function, and protection against light-induced oxidative stress(20), the localized *TYR* signal likely reflects the early establishment of pigment-associated ocular compartments. *INA*, a neuronal intermediate filament gene, showed specific expression in retinal ganglion cells. Previous studies have reported α-internexin expression in developing retinal neuronal lineages, including ganglion cell processes, supporting its association with retinal neuronal differentiation and axonal maturation(21). In addition, *CRYAA* showed strong and highly localized expression in the lens region, especially at later stages. This pattern is consistent with the well-known role of αA-crystallin as a major lens structural protein and molecular chaperone, which helps maintain lens transparency and cellular homeostasis(22). Taken together, the spatially restricted expression of *TYR, INA*, and *CRYAA* captures several key aspects of eye development, including pigmentation, lens formation, and retinal neuronal maturation.

Taken together, these spatial expression patterns demonstrate that HESTA preserves the anatomical context of human embryonic transcriptomes and enables intuitive visualization of gene activity across developmental stages. The observed enrichment of mesodermal, smooth muscle, hepatic, intestinal, neural, and ocular markers is consistent with known developmental biology, while also highlighting the value of HESTA as a reusable spatial transcriptomic resource for studying human organogenesis.

### Re-use Potential

#### Overview of HESTA

HESTA supports the reuse of the human early organogenesis atlas through a web-based interface that connects atlas-level annotations, processed h5ad files, derived matrices and representative visualization modules (https://db.cngb.org/hesta/, Figure 1). The web resource is organized into several functional sections: Home, Spatial Explorer, Single-cell, Organ Atlas, Analysis, Download and Help. Together, these sections provide a structured route from data discovery to spatial visualization, organ-centred interpretation, gene-level query, cross-species comparison, allele-specific expression inspection and data download.

The Home page introduces the scope of HESTA and provides entry points to major data resources and exploration modules. The Spatial Explorer is the main spatial browsing interface, integrating Embryo Overview, Spatial Clustering, Substructure Clustering and 3D Reconstruction. The Organ Atlas module supports organ-centred exploration of selected gene, pathway and regulon data products. The Single-cell module provides cell-type-level reference information for interpreting spatial signals. The Analysis module includes Gene Overview, Cross-Species Comparison and Allele, enabling users to query genes, compare expression patterns across species and inspect spatial allele-specific expression signals. The Download section serve as the data catalogue and download entry point, allowing users to identify available data products, file types, developmental stage coverage, organ or substructure coverage and download links. The Help section provides documentation, tutorials, a data dictionary, h5ad file descriptions and citation guidance.

#### Spatial exploration

The Spatial Explorer provides the primary route for reusing section-level spatial transcriptomic data. Through Embryo Overview, users can first inspect embryo-level and section-level context, including developmental stage, section identity and the anatomical position of each section within the whole embryo. This is important because the interpretation of two-dimensional sagittal sections requires whole-embryo anatomical context.

After selecting a section, users can enter Spatial Clustering to view organ-level spatial annotations and query gene expression patterns within the selected section. The interface supports spatial expression overlays, allowing users to compare gene expression with organ-level annotations in the same anatomical context. For representative sections with high-resolution annotations, Substructure Clustering provides a finer anatomical layer that allows users to inspect local spatial domains beyond organ-level labels.

The 3D Reconstruction module is also part of the Spatial Explorer. It provides a representative three-dimensional view generated from selected consecutive sections, helping users interpret spatial relationships that are difficult to infer from isolated two-dimensional sections. Users can rotate and inspect the reconstructed embryo to understand the relative positions of organs and tissue domains. Because the current 3D resource is based on selected section series rather than all embryos or stages, it should be interpreted as a representative spatial reconstruction resource rather than a complete 3D reconstruction of the entire atlas.

This workflow enables users to move from whole-embryo context to section-level spatial patterns, substructure-level annotations and selected three-dimensional anatomical context. For example, a researcher interested in neural development can locate relevant spinal cord or brain sections, query marker genes, and compare their expression with organ or substructure annotations.

#### Organ-centered reuse of gene, pathway and regulon data

The Organs module supports organ-centred reuse of selected derived data products. It organizes available organ-level data into three complementary layers: gene expression, pathway activity and regulon activity. The gene layer allows users to inspect spatial expression patterns of individual genes within selected organs or anatomical regions. The pathway layer summarizes Gene Ontology Biological Process activity at the spatial bin level, enabling users to explore functional programs beyond individual genes. The regulon layer provides AUCell enrichment scores and supports the interpretation of candidate regulatory programs in spatial context.

These organ-level data products are useful for researchers who want to focus on a specific organ system across developmental stages. For example, users studying heart development can inspect the spatial distribution of cardiac marker genes, compare pathway activity across developmental stages and examine candidate regulatory programs associated with cardiac substructures. Users studying brain, kidney, eye or skeletal muscle development can apply a similar workflow to investigate organ-specific differentiation and spatial regulatory patterns.

#### The analysis section

The Analysis section provides three focused modules for gene-centered exploration, cross-species comparison and allele-specific expression inspection: Gene Overview, Cross-Species Comparison and Allele. These modules are designed to help users start from a gene or a specialized biological question and then connect HESTA to relevant spatial, single-cell, cross-species or allele-specific information.

The Gene Overview module provides a gene-centered entry point for exploring spatial expression patterns across HESTA and related human/mouse embryo datasets available through STOmicsDB(23). Users can select a species and query a target gene using either a gene symbol or gene identifier. The module then retrieves matching spatial transcriptomic datasets and displays gene expression heatmaps across available embryo sections, allowing users to rapidly compare where the queried gene is expressed across human or mouse embryonic spatial datasets. Users can click an individual heatmap or section card to enter the corresponding spatial section viewer.

The Cross-Species Comparison module supports the comparison of gene expression patterns across selected human and mouse developmental datasets. In the current release, this module includes four datasets selected to bridge spatial mapping and continuous developmental trajectories: MOSTA: Mouse Organogenesis Spatiotemporal Transcriptomic Atlas (E9.5–E16.5) (24); HESTA: Human Embryogenesis Spatiotemporal Transcriptomic Atlas (CS12–CS23) (8); The Single-cell transcriptional landscape of human brain development (CS10–CS20) (25); and A single-cell time-lapse of mouse prenatal brain development from gastrula to birth (E8.5–E17.5) (26). While HESTA and MOSTA enable direct cross-species spatial comparisons, the single-cell datasets expand the temporal scope to cover earlier and continuous developmental windows.

Together, this integration allows users to assess whether target genes show conserved or divergent expression across species, platforms, and developmental time.

The Allele module provides precomputed spatial maps for selected genes with significant allele-specific expression. Users first select a tissue section, after which the interface displays the corresponding list of significant genes available for that section. When a gene is selected, HESTA renders a precomputed spatial map showing the anatomical distribution of allele-specific expression signals. This allows users to evaluate whether allelic imbalance is localized to particular organs, substructures or developmental regions without reprocessing genotype, phasing and spatial expression data locally.

Together, these Analysis modules extend HESTA from a section-level spatial browser into a reusable resource for gene-centered, cross-species and allele-specific exploration.

## Data Availability

The HESTA database is freely available at https://db.cngb.org/hesta/.

## Declarations

### Ethics approval and consent to participate

Ethics approval for the human embryonic samples used in this study was obtained by the original atlas study as detailed in the accompanying paper(8), All procedures complied with the Helsinki declaration (2013) and relevant national regulations.

### Consent for publication

This study utilizes publicly available, de-identified datasets originally reported by Pan et al. (8). The original study obtained ethical approvals from the Ethics Committee of Obstetrics and Gynecology Hospital, Fudan University (2021-121) and the Institutional Review Board of BGI-Shenzhen (BGI-IRB 22058). Written informed consent for research purposes was obtained from all participants prior to tissue collection, with additional verbal reconfirmation post-termination. Furthermore, the original authors confirmed that all procedures complied with the “Interim Measures for the Administration of Human Genetic Resources” administered by the Chinese Ministry of Health. As the data used in this study are fully anonymized and publicly accessible, no additional consent for publication is required.

### Authors’ contributions

Z.X. conceived the idea. Y.Z. and L.F. curated the data. T.Y., W.D. developed the web interface. J.C. implemented data visualization, Z.X., W.W. and Y.L. wrote the first draft of the manuscript. Y. G., K.M. and Y. L revised and finalized the manuscript.

## Acknowledgements

Not applicable

